# Timing of increased temperature sensitivity coincides with nervous system development in winter moth embryos

**DOI:** 10.1101/2021.03.16.435649

**Authors:** Natalie E. van Dis, Maurijn van der Zee, Roelof A. Hut, Bregje Wertheim, Marcel E. Visser

## Abstract

Climate change is rapidly altering the environment and many species will need to genetically adapt their seasonal timing to keep up with these changes. Insect development rate is largely influenced by temperature, but we know little about the mechanisms underlying temperature sensitivity of development. Here we investigate seasonal timing of egg hatching in the winter moth, one of the few species which has been found to genetically adapt to climate change, likely through selection on temperature sensitivity of egg development rate. To study when during development winter moth embryos are most sensitive to changes in ambient temperature, we gave eggs an increase or decrease in temperature at different moments during their development. We measured their developmental progression and timing of egg hatching, and used fluorescence microscopy to construct a timeline of embryonic development for the winter moth. We found that egg development rate responded more strongly to temperature once embryos were in the fully extended germband stage. This is the phylotypic stage at which all insect embryos have developed a rudimentary nervous system. Furthermore, at this stage timing of ecdysone signaling determines developmental progression, which could act as an environment dependent gateway. Intriguingly, this may suggest that, from the phylotypic stage onward, insect embryos can start to integrate internal and environmental stimuli to actively regulate important developmental processes. As we found evidence that there is genetic variation for temperature sensitivity of egg development rate in our study population, such regulation could be a target of selection imposed by climate change.

## Introduction

One of the most pervasive and consistent temperature-related impacts of climate change is the advancement of seasonal timing. Between 1950 and 2000 alone, spring phenology advanced for all major species groups by on average 5.1 days per decade (Root et al., 2003). Often, not all species within a food chain shift their seasonal timing at the same rate (Kharouba et al., 2018). As a consequence, there is increased selection on timing through the occurrence of phenological mismatches between two interacting species (Visser et al., 2019). In the face of increased selection, the speed with which species can genetically adapt their seasonal timing will determine their capacity to keep up with climate change (Gienapp et al., 2014; Visser, 2008).

To determine how populations can respond to increased selection on seasonal timing, we need to gain insight into the underlying mechanisms of adaptation to climate change (Visser, 2008). So far, only a few examples of rapid genetic adaptation to climate change have been uncovered (Scheffers et al., 2016), such as later onset of diapause in the pitcher plant mosquito, *Wyeomyia smithii* (Bradshaw et al., 2001), earlier onset of flowering in *Brassica rapa* (Franks et al., 2007), and later timing of egg hatching in the winter moth, *Operophtera brumata* (van Asch et al., 2013). Yet little is known about the genetic basis that allowed for such rapid adaptation of phenological traits (Franks et al., 2012).

Seasonal timing is a plastic trait, allowing species to respond to the large variation in environmental conditions from year to year in order to time key life-cycle events to when conditions are favorable (Hut et al., 2011). For spring feeding insects, it is crucial to time their emergence to the phenology of their host plant, as emerging too early will result in starvation, while emerging too late decreases the nutritional value of their food source (van Asch & Visser, 2007). This is especially important for winter moths, which have only a single generation per year. Adults emerge and lay eggs in winter, which need to hatch in early spring for larvae to feed on young leaves until pupation after four to six weeks (Salis et al., 2017). However, warmer winters advanced winter moth timing of egg hatching more than the timing of budburst of their host tree, pedunculate oak (*Quercus robur*). The resulting phenological mismatch of up to 15 days increased the selection for later timing of hatching, driving the rapid genetic adaptation of the winter moth (van Asch et al., 2013).

Winter moth egg hatching is now better timed to oak budburst despite increasingly warmer winters as eggs hatch later for a given temperature compared to 10 years before (van Asch et al., 2013). To investigate the genetic basis of the rapid adaptation of egg development to temperature, we need to know which components of the underlying mechanism were targeted by selection. As insects are ectotherms, their development rate speeds up with higher temperatures, whereas lower temperatures may constrain development rate (Nedved, 2009). Temperature therefore directly influences timing of development completion (Beldade et al., 2011). However, many insects may be able to regulate the extent to, or the time window in which the environment can affect their development. One well-known mechanism is diapause, an epigenetically programmed developmental arrest that allows insects to regulate the time window when they are most sensitive to changes in ambient temperature (Denlinger, 2002).

Previous work has shown that temperature sensitivity of winter moth eggs varies over the course of development. While timing of egg hatching is affected by temperature fluctuations during the entire egg development period, temperature has a larger impact later in development (Salis et al., 2016). This change in temperature sensitivity indicates that winter moths are especially sensitive to temperature during a specific time window, which forms a likely target for selection by climate change. However, it is unclear when during embryonic development this increased temperature sensitivity occurs.

Here, we determined at which embryonic stage winter moth egg development rate is most sensitive to temperature changes. In two split-brood experiments, eggs were given a two week increase or decrease in temperature at different moments during development, and subsequent developmental progression and timing of egg hatching were measured. Using fluorescence microscopy, we constructed a timeline of embryonic development for the winter moth and tested in which development stages egg development rate responded most strongly to temperature increases or decreases. From previous work, we expected that temperature affects egg development rate at every embryonic stage, but with larger effect sizes at later stages. Knowing at which stages embryos are most sensitive to their environment will be instrumental to determine potential targets of selection to explain the rapid genetic adaptation to climate change in the winter moth.

## Methods

We conducted two split-brood experiments to determine the effect of temperature on winter moth egg developmental rate, and whether this effect changes over the course of development (following Salis et al., 2016). We collected eggs in 2018 and 2019 from wild winter moth females caught during the peak of adult emergence in a forest in Doorwerth, the Netherlands (Catch dates: November 26 and 29, and December 3 2018; November 25, 28 and December 2, 2019). At the start of each experiment (December 14, 2018 and December 13, 2019), clutches (ranging from 45 to 191 eggs) were placed in climate cabinets set at a constant baseline temperature of 10°C. Then from the second week onwards, every week four clutches received a two-week temperature treatment. In 2018-2019, eggs received treatment in weeks 2-8 (28 clutches), and in 2019-2020 eggs received treatment in weeks 2-13 (48 clutches). Clutches were sequentially assigned over treatment weeks such that the catch dates were spread evenly across experimental groups.

In treatment weeks, each clutch was divided into 4-7 sub-clutches of preferably 25 eggs, with at least 15 eggs. One sub-clutch was sampled before the start of the temperature treatment. The remaining sub-clutches were divided over three treatments, transferred to either a warmer (15°C) or a colder treatment (5°C), or remained at baseline temperature (10°C). After two weeks of treatment, eggs were either placed back at 10°C to record timing of hatching (2019-2020), or they were sampled to measure the direct effect of temperature changes on developmental progression (2018-2019: weeks 2-8 and 2019-2020: weeks 9-13). Sampled eggs were dechorionated with 50% bleach, fixated with 4% formaldehyde, and dehydrated gradually in methanol (protocol adapted from Brakefield et al., 2009). After storage in 100% methanol at −20°C, whole eggs were then gradually rehydrated and imaged with fluorescence microscopy to determine the development stages of the embryos, using 4′6’-diamidino-2-phenylindole (DAPI) staining which binds to DNA.

In 2018-2019, an additional five clutches were kept at 10°C until hatching to check the total duration of development at this temperature. In 2019-2020, an additional five clutches were sampled regularly from one week before the start of the experiment until the start of the treatments in week 2 to define early development stages.

### Statistical analysis

All statistical analyses were performed using R v. 3.6 (R Core Team, 2019). To test for the effects of temperature treatment on development rate, we used mixed models in a Bayesian framework. For the effect on timing of egg hatching (the ‘hatching dataset’), we used a linear mixed model with the observed hatching date for each embryo in April days as response variable. For the direct effect on developmental progression (the ‘imaging dataset’), we used an ordinal mixed model with the observed development stage for each embryo that was imaged as response variable. The development stages were scored in arbitrary categories, chosen because they could be readily distinguished by microscopy. Because we only know the order and direction of development for these categories, a continuation ratio ordinal model was used for which Pr(Y>i|Y≥i) (Harrell, 2015). This gives the probability in log odds of falling into a higher level than the one observed, given that an embryo can only stay in a particular development stage or continue to the next stages. This model does not make any assumptions about the absolute distance between development stages. We used the R package *brms* (Bürkner, 2017) to fit both models with random effects.

For both models, we used weakly informative normal priors for both intercepts and fixed effects (mean=0, SD=10) to initialize the models (Gelman et al., 2017). We included temperature treatment and treatment week as fixed effects, as well as the interaction between the two. Treatment week was included as a factor, as we are interested in the differences in treatment effects between weeks. Including such group-level predictors addresses the multiple comparisons problem in Bayesian analysis (Gelman et al., 2012). As covariates, we included female catch site and date. Catch tree was included as a random effect, as winter moths can show local adaptation (Dongen et al., 1997). We also included a random intercept for clutch as well as a random slope for treatment per clutch, as the winter moth’s genetic adaptation to climate change suggests genetic variation in both baseline development speed and temperature sensitivity. Removing the covariates and the tree the female was caught on as random effect did not diminish model fit (Watanabe–Akaike information criterion expected log pointwise predictive density difference (WAIC elpd_diff)=+6.4, SE=2.6 and WAIC elpd_diff=+0.8, SE=0.2) nor did it affect the estimates for temperature treatment and treatment week (Fig. S1 and S2). Therefore, we decided to use these more parsimonious models as our final models. Posteriors for all model parameters converged (R_hat_=1.00) with effective sample sizes of >2000 (Table S1-2).

As the effect of temperature on development speed in insect embryology is well established to be directional (Nedved, 2009), we used one-tailed tests at a significance level of α=0.05. To test our hypothesis that differences in development rate between warm and cold treatments are present after every treatment period, we compared treatments within each treatment week. To determine when the effect of temperature on winter moth egg developmental rate changes over the course of development, we compared the effect size of the warm and the cold treatments relative to the constant baseline between treatment weeks.

## Results

### Timeline of winter moth embryonic development

Given the weekly sampling of eggs, we constructed a timeline for winter moth embryonic development at a constant 10°C. We used the timeline of a related species from the same Geometridae subfamily as the winter moth as guidance (Wall, 1973) and defined 20 development stages, which were easily distinguishable with whole-egg fluorescence microscopy using DAPI staining (Figure 1). Recently laid eggs in stage 1 were still green, but turned orange over the course of a week. On average, embryos took approximately 14 weeks at a constant 10°C to complete embryonic development (Fig. S3).

**Figure 1.**
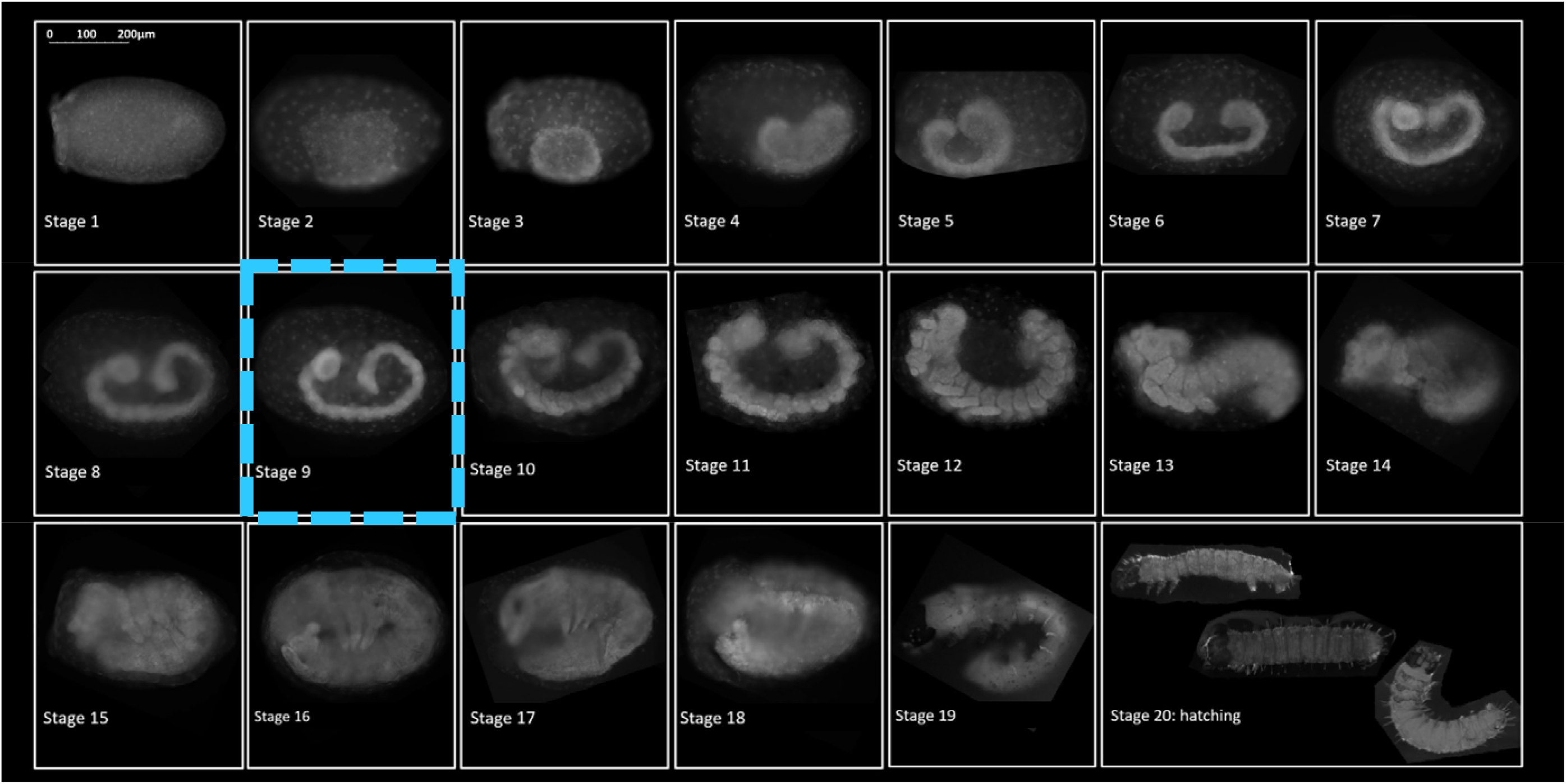
Timeline of winter moth embryonic development. We identified 20 distinct development stages in the winter moth, similar to the embryonic development timeline of a related Lepidoptera species from the same Geometridae subfamily (Wall, 1973). The fluorescent microscopy images shown are typical representations of each development stage. See main text for a detailed description. In our experiments, we observed an increase in egg temperature sensitivity after embryos had reached stage 9 in which they finish segmentation and have formed a rudimentary nervous system.

Figure 1 depicts a typical image for each of the 20 development stages we identified for winter moth embryonic development. The blastoderm stage was defined as stage 1. At stage 2, the orange-pigmented serosa migrated over the germ rudiment, evidenced by the large serosal nuclei overlying the denser cells of the germ rudiment. This germ rudiment further condensed into a cup shape (stage 3), although not as extremely as observed in *Chesias legatella* (Wall, 1973), and at the borders of the germ rudiment a thicker rim of amniotic cells formed (Gaumont, 1950). As the embryos started to elongate into a germband, the head lobes started to form (stage 4), and the formation of both head and tail pouches (Wall, 1973) became prominently visible in stage 5. Subsequently, the germ band sunk deep into the yolk and the head and tail pouches reduced in size (stage 6). As embryos elongated further, head and tail nearly touched each other (stage 7), but no constrictions in the germ band were visible, until segmentation of the anterior segments started (stage 8). As segmentation continued towards the tail and completed (stage 9), the germband reached its maximum length, and thoracic segmentation became more refined. At this stage, the brain, central nerve chord, and abdominal ganglia have formed, according to Gaumont (1950). In stage 10, head and thorax appendages started to arise, with embryos still having a relatively thin posterior abdomen. The head appendages then became more rod shaped and started to fuse together (stage 11), while the thoracic legs grew longer, and the posterior abdomen thicker. At stage 12, we observed germband retraction, with embryos in a C-shape position and the head parts almost completely fused together. Then the tail moved away from the head until embryos flipped their tails towards the ventral side at the start of revolution (stage 13: katatrepsis, Panfilio, 2008). Embryos elongated further with the tail moving towards the thorax (stage 14), until they were completely in a U-shape (stage 15). The back of the head smoothed out, and the mouth became directed downwards, while embryos increased in length (stage 16) and we started observing a clasper at the end of the tail. Pigmentation started first at the eye and jaw (stage 17), and where before embryos had had an open back, from this point forward we observed the progression of dorsal closure. As pigmentation continued, DAPI penetration reduced, and pigmentation showed as black areas that did not reflect light. A black cap formed on the head of the embryos, and sclerotization of the body started (stage 18). In this stage, embryos went through a final elongation with the head tucked in towards the center of the egg. With pigmentation completed (stage 19), fully grown caterpillars could be observed with a light microscope lying in a transparent chorion, which always burst during the fixation process. The last stage (stage 20) we defined as the moment of egg hatching.

Ultimately, we are interested in whether the effect of temperature on development rate changes during development. To aid in the interpretation of the direct effect of temperature on developmental progression and to be able to compare it to the effect on timing of hatching, we linearized the development timeline at a constant 10°C with a locally estimated scatter plot smoothing (loess) model. This allowed us to translate the observed development stages into time units, expressed as the number of days at a constant 10°C (Fig. S3).

### Temperature effect on egg development rate

In both experimental years, egg development rate responded more strongly to temperature once embryos had passed stage 9, in which they finish segmentation (Figure 1 and 2). We observed this change in temperature sensitivity in response to two weeks of temperature treatment both in developmental progression (Figure 2A) and at timing of hatching (Figure 2B).

**Figure 2.**
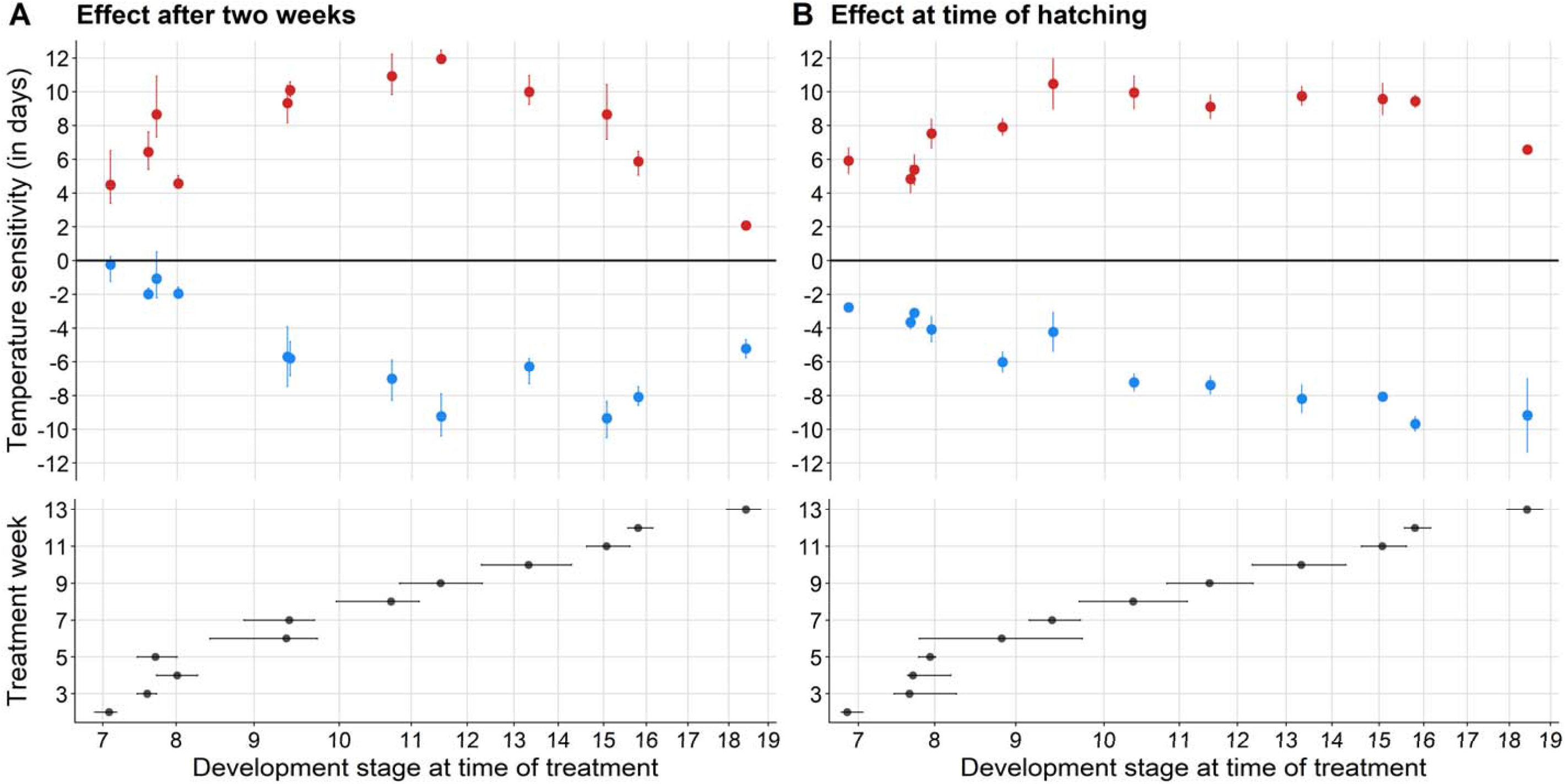
Change in winter moth temperature sensitivity during development, measured (A) directly after a two-week temperature treatment as development progresses and (B) at timing of hatching. Temperature sensitivity is expressed in number of days embryos were delayed (blue) or advanced (red) in response to a two-week temperature treatment compared to development at a constant 10°C (zero line), with for (A) medians ±IQR and for (B) means ±SE. Temperature treatment consisted of two weeks at 5°C (blue) or 15°C (red) at different moments during development. Lower panels show the median observed development stage ±IQR at the start of a treatment for each experiment. X-axis spacing reflects the relative timing of each development stage at a constant 10°C (Fig.S3). All points have been adjusted for between-clutch variation (A: N=28 + 48 clutches; B: N=48 clutches). To aid interpretation, effect sizes for developmental progression (A) have been translated from the observed discrete development stages (Fig.S4) to time units, expressed as the number of days at a constant 10°C, with a loess model (Fig.S3). For both datasets, comparing effect sizes for the 5°C and 15°C treatments between timepoints shows an increase in temperature sensitivity after embryos have reached stage 9 in which they finish segmentation (Table S5-8).

For developmental progression, we found that in every treatment week, embryos from each treatment group progressed in development compared to the development stage observed before treatment (Table 1: estimated mean probabilities are all positive log odds, Fig. S4). The probability of observing a later stage of development was always significantly higher for embryos in the warm treatment compared to the cold and baseline treatments after two weeks (Table 1: 15 vs. 5°C, P<0.05, Table S3). Thus, eggs of the warm treatment were always significantly further along in development. When we compared the cold treatment to the constant baseline, we only observed a significant delay in development from treatment week 6 onwards (Table 1: 5 vs. 10°C, P<0.05, Table S3), when embryos received treatment after they had passed stage 9: the completion of segmentation (Figure 1). At the time of segmentation, the effect size of temperature treatment significantly increased when compared between weeks (Table 1, Fig. S5A and S6-7, Table S5-6, P<0.05). When we translated the effect size in each week to number of days at 10°C (Fig. S3), we observed that a warm treatment administered after segmentation led to an advance of 9-12 days compared to development at a constant 10°C, while this advance was only 4-6 days before segmentation (Figure 2A). An increase in the effect size of the cold treatment also became apparent at this stage: once embryos had finished segmentation a cold treatment of two weeks resulted in a delay of 6-10 days compared to only 0-2 days before (Figure 2A).

**Table 1.**
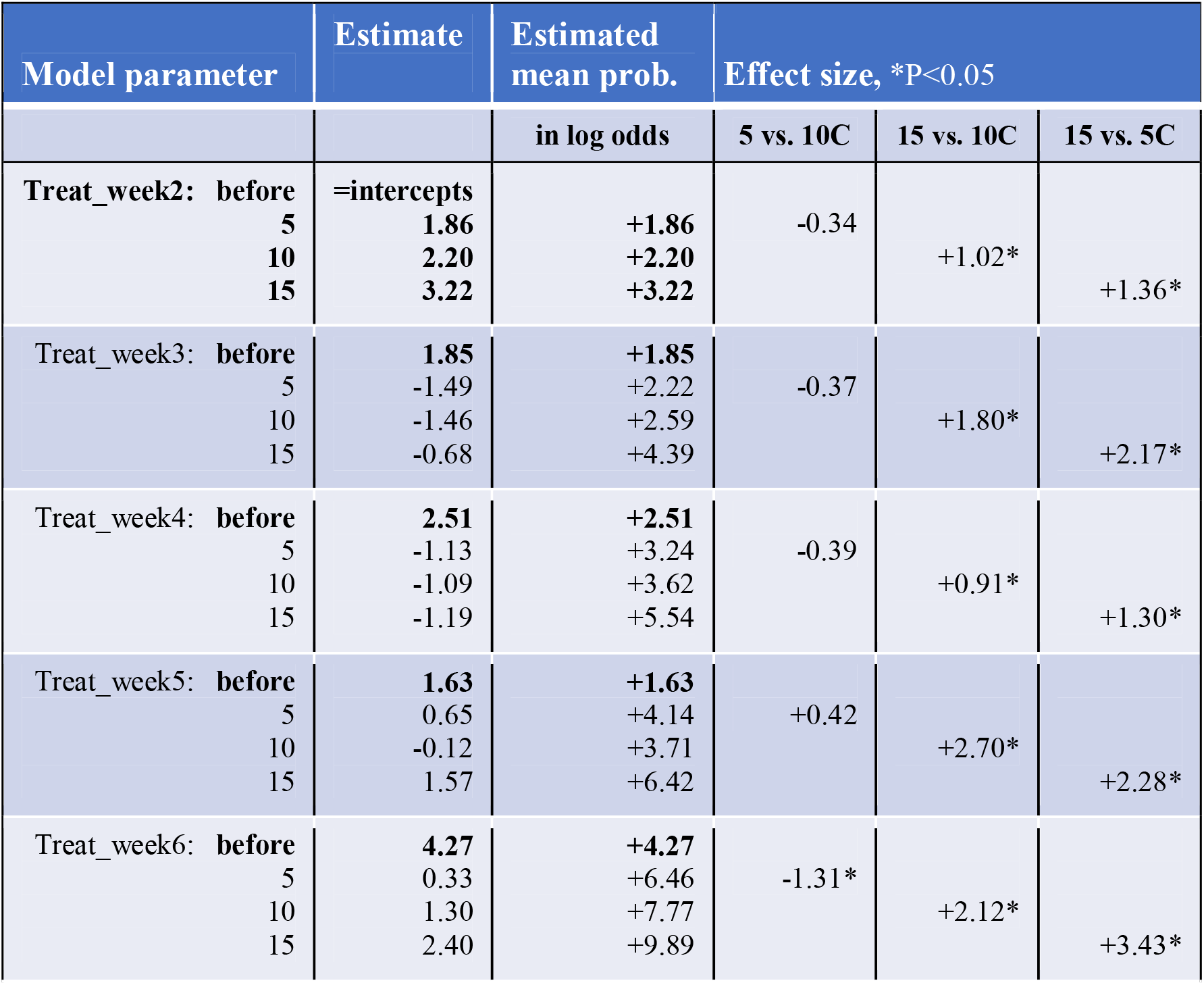
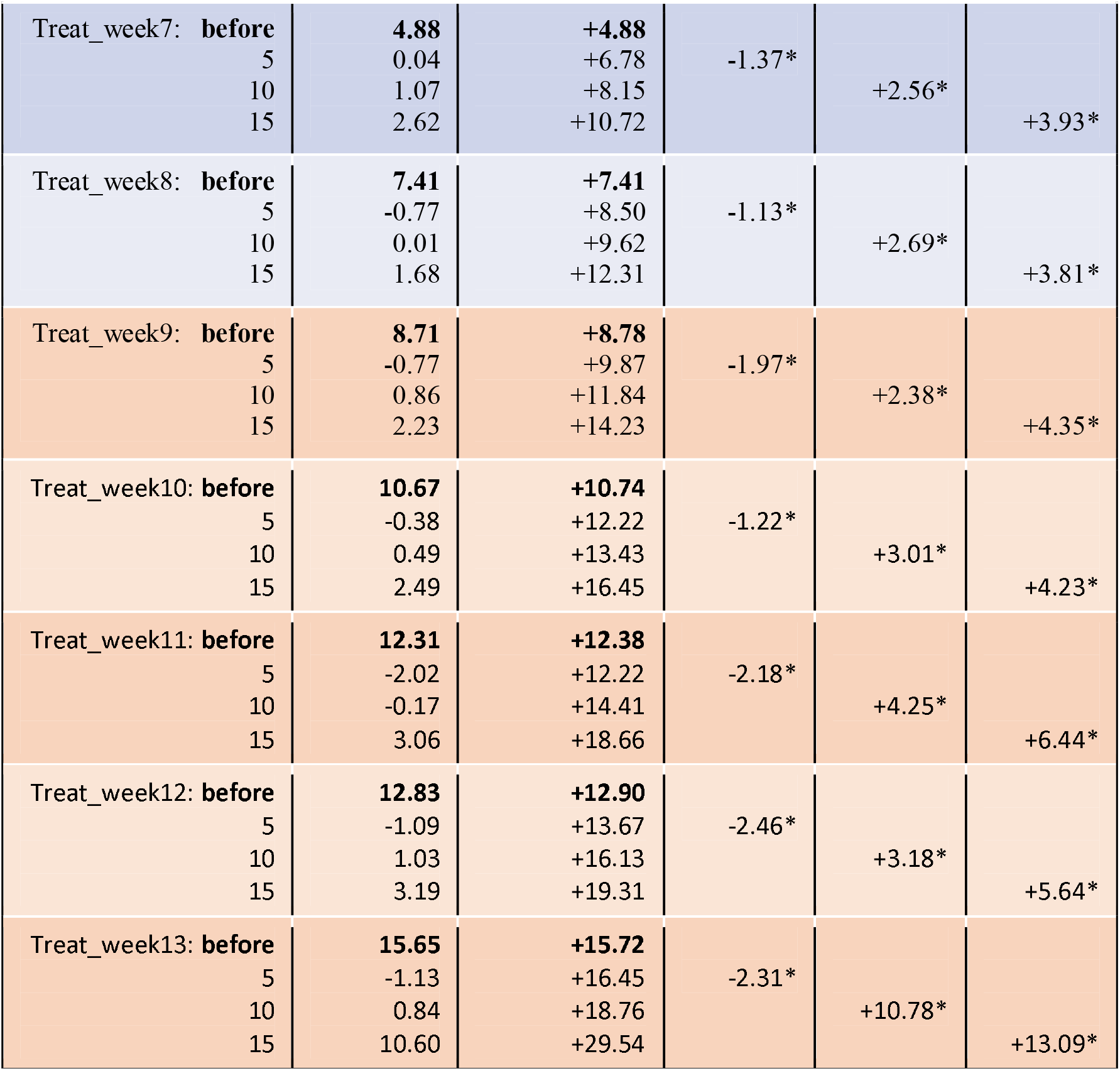
Model output and effect sizes for temperature effect on developmental progression. Estimates are expressed in log odds. Estimated mean probabilities and effect sizes are expressed as change in log odds, with reference levels in bold. In 2018-2019, treatments were given in weeks 2-8 from the start of the experiment (blue rows, N=28 clutches). In 2019-2020, eggs were sampled weekly (=before), but treatment was only administered in weeks 9-13 (orange rows, N=48 clutches). As we observed winter moth embryos from 18 different developmental stages in the experiment (stage 2, 3, 5-20), the model includes 17 intercepts that denote the thresholds between these developmental stages. Asterisks denote significant within-week comparisons. See for full model output Tables S1 and S3.

A similar shift in temperature sensitivity was observed at timing of hatching (Figure 2B). All treatments significantly differed from each other regardless of the moment at which temperature treatment was administered during development (Table 2: effect size, P<0.05, Table S4), confirming that winter moth embryonic developmental rate is sensitive to temperature during the entire egg stage. Embryos that received a warm treatment always hatched earlier compared to development at a constant 10°C and to the cold treatment (Table 2: 15 vs. 10°C and 15 vs. 5°C, negative effect sizes), while embryos that received a cold treatment always hatched later (Table 2: 5 vs. 10°C positive effect sizes). However, the magnitude of the temperature effect on timing of hatching changed over the course of development. The effect size of temperature treatment in the weeks after which embryos had finished segmentation significantly increased compared to the weeks before (Table 2, Fig. S5B and S8-9, Table S7-8, P<0.05). For the warm treatment, embryos that were moved to 15°C when they had passed stage 9 were 8-10 days advanced compared to hatching at a constant 10°C, while they were only 5-8 days advanced when they were moved to 15°C earlier in development (Figure 2B). Similarly, the largest delay in hatching after a cold treatment was observed for embryos that were moved to 5°C after they passed stage 9, going from a 3-6 days delay to a 7-10 days delay compared to hatching at a constant 10°C (Figure 2B).

**Table 2.**
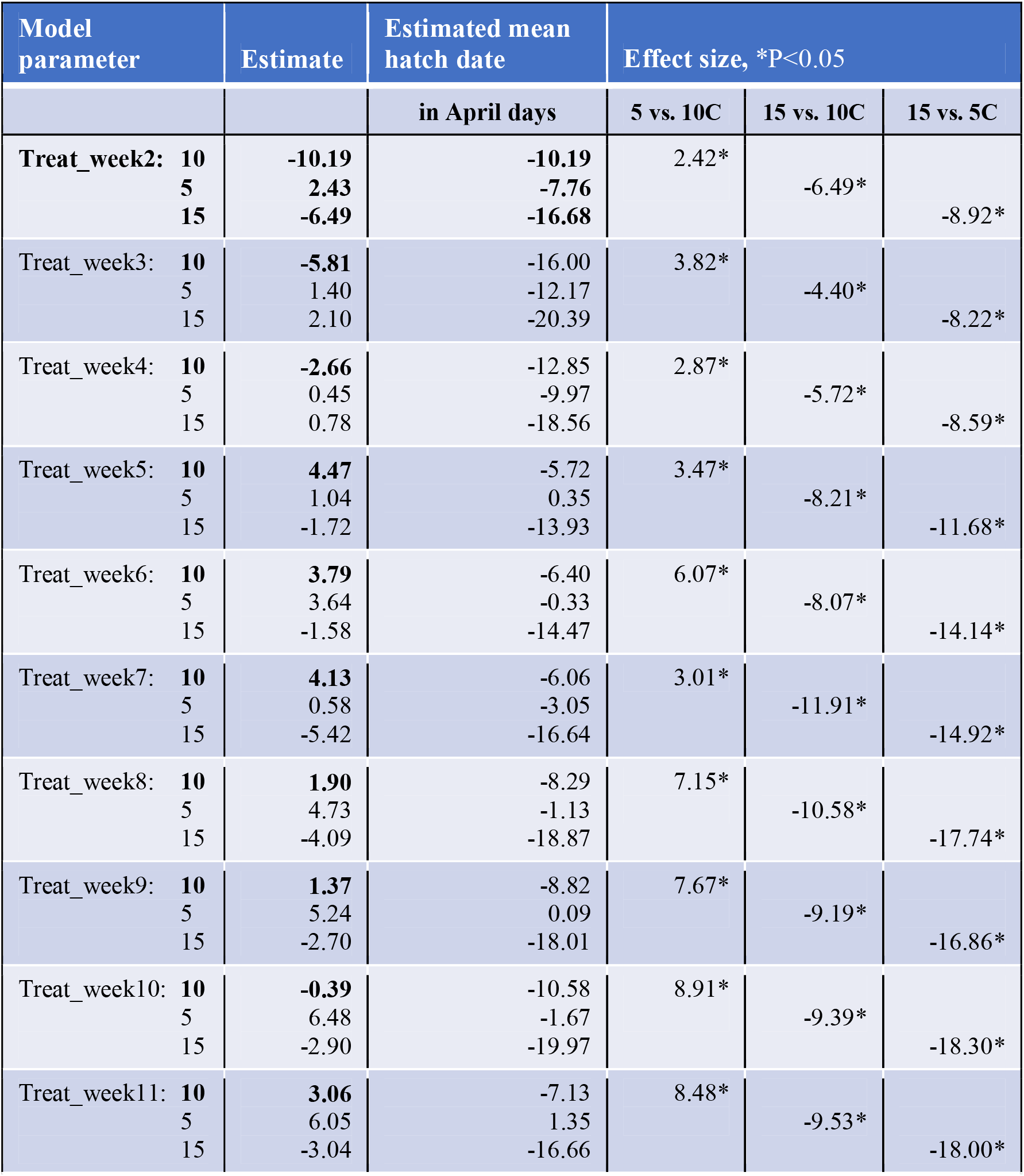
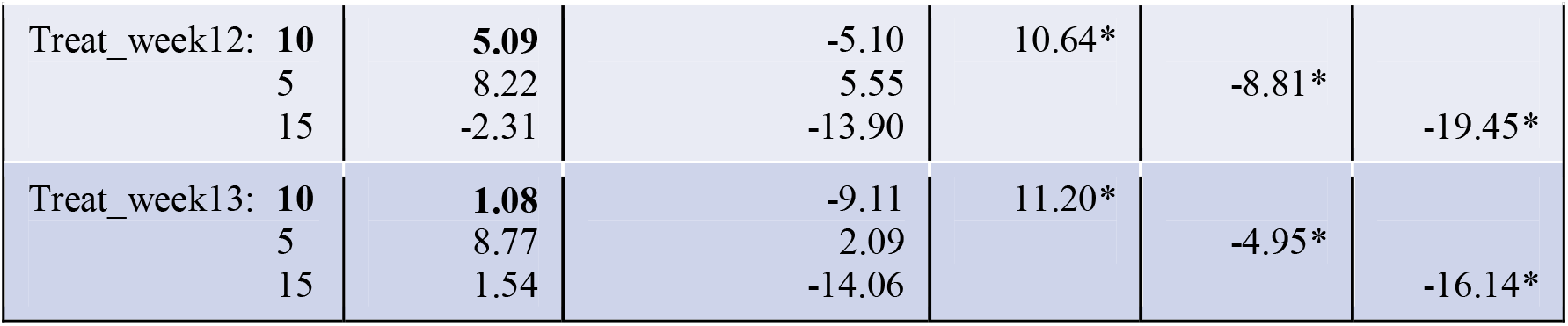
Model output and effect sizes for temperature effect on timing of hatching. Estimates and estimated means are expressed in April days, with reference levels in bold. Negative estimated means indicate that clutches hatched before April 1^st^. Effect sizes are expressed in days, with negative numbers meaning an advance in timing and positive numbers a delay. Asterisks denote significant within-week comparisons. See for full model output Tables S2 and S4.

### Variation in development speed and temperature sensitivity

There was high between-clutch variation in development speed. At a constant 10°C, the earliest clutch and the latest clutch hatched 18 days apart (mean=April day −9.71, SD=8.07). Moreover, there was high within-clutch variation with on average an IQR of 7.34 days within-clutch (SD=3.73). This high variation was also visible in the range of different development stages observed at each time point (Fig. S3).

The high variation in hatch dates and development stages could not solely be explained by the temperature environment. The random intercept for clutch as well as the random slope for treatment per clutch were significantly different from zero in both models of egg development rate (Table S1 and S2, P<0.05). This means that both baseline development speed and temperature sensitivity depended on clutch and probably had a genetic basis.

## Discussion

Temperature sensitivity of winter moth egg development rate was previously found to change over the course of development. The mechanism behind this change in temperature sensitivity represents a potential target of selection on seasonal timing imposed by climate change. To gain insight into the underlying mechanism, we investigated at which embryonic stage winter moth egg development rate is most sensitive to changes in temperature. We found a switch from weak to strong temperature sensitivity once embryos had finished segmentation and were in the fully extended germband stage.

As ectotherms, insect development rate is largely dependent on ambient temperature (Nedved, 2009). This is also reflected in our results: embryos that had received a warm treatment for two weeks were always advanced in development and hatched earlier, while embryos that received a cold treatment were always delayed compared to the control. This seems to suggest that winter moth embryos do not have egg diapause. Interestingly, winter moth embryos did condense into a cup-shape, which resembles the pyriform embryonic stage observed in many Lepidopteran species with egg diapause (Behrens, 2012). Indeed, in *C. legatella* embryos enter diapause in this cup-shaped stage (Wall, 1973). However, the condensation was less extreme in the winter moth and embryos had formed a germband within two weeks at a constant 10°C. In contrast, diapausing *C. legatella* embryos go through a period of stasis before germband development resumes after a prolonged period of cool temperatures (Wall, 1974).

The extent to which winter moth development rate was affected by changes in temperature shifted over the course of development, as previously found by Salis et al. (2016). Our results indicate that winter moth embryonic development can be divided into two phases of temperature sensitivity. In both experiments, the switch from weak to strong temperature sensitivity occurred once embryos were in the fully extended germband stage. The switch seems to have occurred progressively rather than abruptly, with a strong increase in sensitivity over the course of two to three weeks, followed by a gradual approach towards a maximum advancement or delay of 10-12 days, which is close to the two-week treatment duration we used. This graduality may either reflect the underlying regulating mechanism of temperature sensitivity or it may be due to the large variation in development rate both within and between clutches.

The fully extended germband stage, where we observed the switch from weak to strong temperature sensitivity, coincides with two developmental events. Firstly, it coincides with the development of a rudimentary nervous system in the winter moth (Gaumont, 1950). Interestingly, this is the phylotypic stage at which all insect embryos resemble each other and have developed a rudimentary nervous system (Sander, 1983; Slack, 2003). This represents the intriguing possibility that insect embryos can start to integrate internal and environmental stimuli to actively regulate important developmental processes. An important aspect for such regulation might be the development of thermosensory neurons, allowing embryos to start sensing ambient temperatures apart from the direct effects of temperature on enzyme kinetics. For example in *Drosophila*, mutants that lack thermosensory neurons are unable to behaviorally respond to changes in temperature, which implies the involvement of cognitive control (Soto-Padilla et al., 2018).

The second major developmental event in the fully extended germband phase is a peak in the hormone ecdysone, as has been shown in *Drosophila* (Kozlova et al., 2003). Ecdysone is a key life-history hormone well known for its regulatory role in timing of insect metamorphosis (Adams, 2009). For example, diapause termination involves an increase in sensitivity to ecdysteroids by the upregulation of ecdysone receptors (Denlinger, 2002) and ecdysone temporal expression also seems to play an essential role in insect embryonic development (Buszczak et al., 1999). If the temporal pattern of ecdysone signaling is dependent on the environment, this signaling could act as a gateway during development as it does in the developmental plasticity of *Bicyclus anyana*. In this species, adult seasonal morphotype was found to depend on ambient temperatures experienced during caterpillar development, with the timing of the peak in ecdysteriod hormones occurring earlier when individuals were placed in warmer temperatures (Oostra et al., 2011).

Rapid climate change results in pervasive changes in local environments, driving shifts in the seasonal timing of many species (Root et al., 2003). This phenotypic plasticity alone is expected not to be sufficient to deal with climate change (Gienapp et al., 2014), as was the case for the winter moth (van Asch et al., 2013). As such, environment dependent regulation of the timing of development represents a likely target of selection in the face of climate change. Gateway mechanisms might be especially important for rapid genetic adaptation. For example in the pitcher plant mosquito, climate change resulted in a genetic shift in the threshold for seasonal timing: critical photoperiods for diapause induction shortened (Bradshaw et al., 2001).

The genetic adaptation of the winter moth to climate change resulted in later egg hatching despite warmer winters (van Asch et al., 2013). Our analysis indicated that both baseline development speed and temperature sensitivity depended on clutch. As the response of egg hatching to temperature was previously found to be highly heritable (*h^2^*=0.63-0.94, van Asch et al., 2007), this likely points to genetic variation present in our study population for these traits. This is in line with van Asch et al. (2013) who find that the winter moth genetically adapted its temperature dependent development rate in response to climate change.

The switch in temperature sensitivity at the time of nervous system development we find here, as well as the presence of genetic variation in temperature sensitivity in our population, can guide future studies on when to look at genes involved in the regulation of developmental timing. We have few examples of species which have been found to genetically adapt to climate change (Scheffers et al., 2016). Characterizing the genetic adaptation in wild populations like the winter moth will help in determining the factors that influence the evolutionary potential of wild insect populations. Knowing the processes and the genes involved in adaptation will be essential for the assessment of vulnerability to climate change. Populations that show genetic variation in genes relevant for climate change adaptation are predicted to be better able to keep up with the high rate of global warming, making them less vulnerable to extinction (Norberg et al., 2012).

## Acknowledgements

We thank B. van Lith, P. de Vries, C. Mateman, and A. Pijl for their technical support and help with lab work during the two experiments.

